# Investigating Enzyme Function by Geometric Matching of Catalytic Motifs

**DOI:** 10.64898/2026.02.10.705182

**Authors:** Raymund E. Hackett, Ioannis G. Riziotis, Martin Larralde, António J. M. Ribeiro, Georg Zeller, Janet M. Thornton

## Abstract

The rapidly growing universe of predicted protein structures offers opportunities for data driven exploration but requires computationally scalable and interpretable tools*(1–3)*. We developed a method to detect catalytic features in protein structures, providing insights into enzyme function and mechanism. A library of 6780 3D coordinate sets describing enzyme catalytic sites, referred to as templates, has been collected from manually curated examples of 762 enzyme catalytic mechanisms described in the Mechanism and Catalytic Site Atlas*(4–6)*. For template searching we optimised the geometric-matching algorithm Jess*(7)*. We implemented RMSD and residue orientation filters to differentiate catalytically informative matches from spurious ones. We validated this approach on a non-redundant set of high quality experimental (n=3751, <40% amino acid identity) enzyme structures with well annotated catalytic sites as well as predicted structures of the human proteome. We show matching catalytic templates solely on structure is more sensitive than sequence- and 3D-structure-based approaches in identifying homology between distantly related enzymes. Since geometric matching does not depend on conserved sequence motifs or even common evolutionary history, we are able to identify examples of structural active site similarity in highly divergent and possibly convergent enzymes*(8)*. Such examples make interesting case studies into the evolution of enzyme function. Though not intended for characterizing substrate-specific binding pockets, the speed and knowledge-driven interpretability of our method make it well suited for expanding enzyme active-site annotation across large predicted proteomes. We provide the method and template library as a Python module, Enzyme Motif Miner, at https://github.com/rayhackett/enzymm.

## Introduction

Nature has evolved many enzyme families, with differences and commonalities in mechanism, catalytic sites and substrate specificity*(9)*. However these exact functional annotations of catalytic sites and specific residues are often lacking, even when the structure has been experimentally determined or predicted. With the proliferation of predicted protein structures increasingly covering major sequence databases*(1–3)*, structural approaches offer new avenues to address this problem. Distantly related proteins may only share local similarities in conserved regions such as enzyme active sites.

Most methods for detecting homology are sequence-based due to their computational efficiency, but become increasingly unreliable below ∼40% sequence identity. Here structural methods have proven more sensitive in identifying cases of divergence between more distantly related proteins. Particularly useful are hierarchical domain-level comparisons such as CATH*(10–13)*. We set our focus on identifying shared arrangements of catalytic residues, which complements fast but coarse structural alignment tools like Foldseek *(14)*. Enzyme active sites are typically highly conserved and the relative position and orientation of residues are constrained by their chemical mechanisms, providing a robust basis for comparisons. This has made searchable templates of atom coordinates a method of choice for detecting such similarities.

Searchable templates were first introduced by Wallace et al.*(15)* and further developed into Jess*(7)*, a geometric matching algorithm which we have optimized for this work. A template is interpreted as a set of constraints with respect to coordinate geometry and atom - or amino acid type. The algorithm will find all residue arrangements in a set of target protein structures sufficiently similar to a provided library of templates. Notably, Laskowski et al. make use of enzyme active site and binding site templates matching using Jess in ProFunc, among other means of detecting protein similarity*(16,17)*. ProFunc relies on SiteSeer scores measuring local similarity around matches based on pairs of similar residue types and their relative sequence position to filter out biologically spurious (false positive) matches at the cost of detecting convergent or more divergent matches. While ProFunc is accurate for single query tasks, it only provides a web interface prohibiting large scale applications. In a different approach by Spriggs et al.*(18)*, residue arrangements are detected by their similarity in functional residue angles (not unlike our own residue orientation angles) in an angle-based graph representation of structures. A further conceptually comparable approach, pyScoMotif, relies on prior indexing of structures into geometric bins, while reducing residue side-chains to averaged positions*(19)*. As this method does not provide prespecified templates, it is geared towards small scale custom searches. Other methods compare predicted physico-chemical properties of protein structure surface representations *(20,21)*.

For detecting catalytic sites in protein structures at scale we developed Enzyme Motif Miner, a knowledge-based approach leveraging templates derived from the Mechanism and Catalytic Site Atlas (M-CSA)*(4,5)*, a database of manually curated and annotated catalytic mechanisms. We hope our approach might find application in areas such as enzyme design, protein clustering, homology detection and elucidation of enzyme evolution as our tool is well positioned to detect distantly related, divergent enzymes. Accordingly, we validate Enzyme Motif Miner on a set of non-redundant structures sharing at most 40% sequence identity from the Protein Data Bank (PDB)*(22,23)*. We further hypothesise that enzymes which have evolved independently over a long time but converged to catalyse a similar reaction on a similar substrate structure may be constrained by the available chemistry of life to share some active site commonality. In the context of paradigms of convergence recently presented by Riziotis et al., such enzymes might share mechanistic and structural similarity*(8)*. To distinguish cases of convergence and divergence, we contrasted template similarity with overall protein fold conservation from CATH annotations and enzyme function by EC annotations*(24)*.

The development of Enzyme Motif Miner was directly motivated for application to predicted protein structures enabled by among others Alphafold*(1,2)*. To exemplify this use case, we carried out analyses on two datasets of predicted protein structures. 1.) Alphafold2 predictions of the Swiss-Prot*(25)* reviewed human proteome and 2.) novel protein folds in the CATH database as identified in the encyclopedia of domains*(26)* from the entire AlphafoldDB(2). With Enzyme Motif Miner we aim to provide insight into the evolution of enzyme function and relationships between distantly divergent enzymes, while facilitating transfer of catalytic and mechanistic annotations to previously uncharacterised structures.

## Results and discussion

### Implementation and availability

For detecting catalytic sites in protein structures at scale we developed Enzyme Motif Miner as a python package, including its library of catalytic templates from the M-CSA, which can be used both through the command line or via a modern Python API (Fig 1A). For template searches, Enzyme Motif Miner depends on PyJess, which we developed from Jess (see methods). We also developed a webserver (https://www.ebi.ac.uk/thornton-srv/m-csa/enzymm/) which allows users to either upload and search a structure of their own or to search deposited structures in the PDB*(22,23)* or AlphafoldDB*(2)* against our library of templates. Matches can be interactively explored for easy and quick analysis.

**Figure 1.**
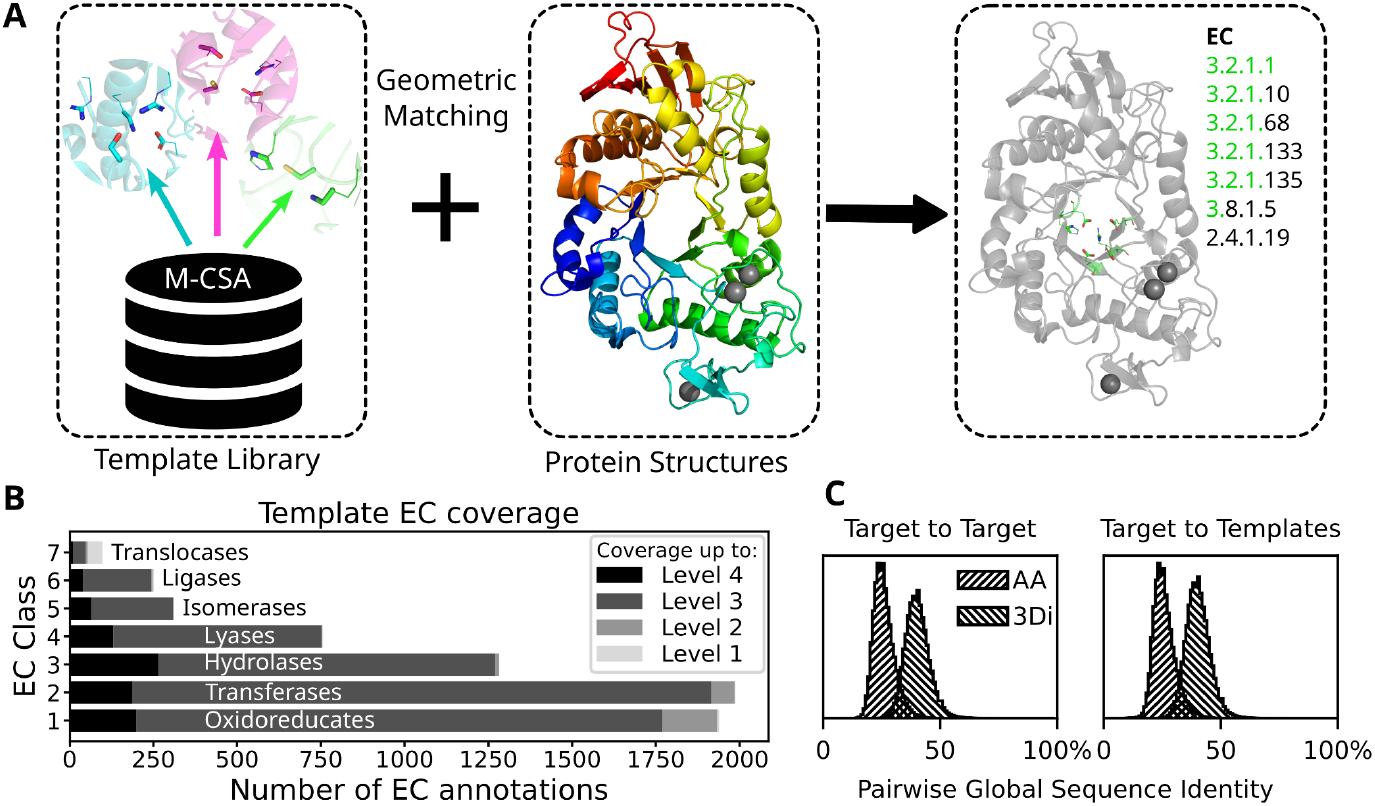
(A) A library of catalytic templates was derived from the Mechanism and Catalytic Site Atlas. Similar catalytic residue patterns can be quickly identified in any protein structure by geometrically matching templates. (B) Our template library provides coverage of nearly all EC numbers listed in KEGG across all major classes up to at least the 3rd level. For example, though our template library does not explicitly include EC number 3.2.2.14, EC numbers of other N-glycosyl hydrolases (3.2.2.) are represented. (C) Evaluation was performed on non-redundant protein structures below 40% amino-acid sequence identity. Distribution of global pairwise sequence identity between target structures or between targets and templates in amino-acid and 3Di sequence space are shown.

Enzyme Motif Miner is computationally efficient. Through various optimizations (see methods) the underlying PyJess algorithm runs ∼12x faster on 12 cores than an earlier threadsafe version of PyJess. On a low-end consumer laptop with 6 cores/threads Enzyme Motif Miner can scan 1000 *E. coli* protein structures in 89s (0.09s per structure) and 1000 human protein structures in 229s (0.23s per structure). Modeling the speedup over parallel threads with Amdahl’s law*(27)* suggests a parallelizable fraction of 86% (Fig. S16), highlighting the scalability of our method on multi-core machines.

### Template selectivity depends on size

We have previously collected recurring catalytic residue arrangements from the M-CSA in 3D*(28,29)*. Structures including residue substitutions were saved in the form of a template. Each template specifies a 3D arrangement of 3 to 8 catalytic residues. Following the approach introduced by Riziotis et al.*(30)* we have constructed a library of 6780 different templates across 762 entries in the M-CSA all derived from experimentally resolved enzyme structures, which comprise 897 complete and 57 partial (e.g. 3.2.1.-) EC numbers. The resulting template library provides coverage of 95% of all EC numbers in KEGG to the third level. This drops to only 13.5% if we include all EC numbers to the 4th level, which include homologous enzymes acting on different substrates that are represented by a single entry in M-CSA, since these often share a conserved mechanism despite different substrates(Fig. 1B)*(6)*.

As each template represents different arrangements of residues, templates inherently differ in specificity. Larger templates with more residues, are much more specific and are expected to occur less frequently by chance, while also being more susceptible to evolutionary divergence or conformational change. We evaluated template matching at different parameters on the validation dataset of 3751 non-redundant (max. 40% pairwise sequence identity, Fig. 1C) experimental protein structures from the PDB with catalytic annotations by homology to reference entries in the M-CSA. Selectivity was assessed across pairwise distance parameters varying between 0.7 Å and 2.0 Å. A match was considered correct if it included at least three annotated catalytic residues.

The number of matches per target increased with progressively higher, more lenient pairwise atom distances as did the total number of targets and templates for which any matches were found (Fig. 2A, Fig. S3-4). The vast majority of matches occurred with 3-residue templates especially as the pairwise distance was relaxed (Fig. 2B). This is explained by smaller residue arrangements being more likely to occur at random than larger patterns. Given the increase in matches for smaller templates at higher pairwise distances, we sought to differentiate correct matches to active sites from false matches originating from non-catalytic parts of protein structures.

**Figure 2.**
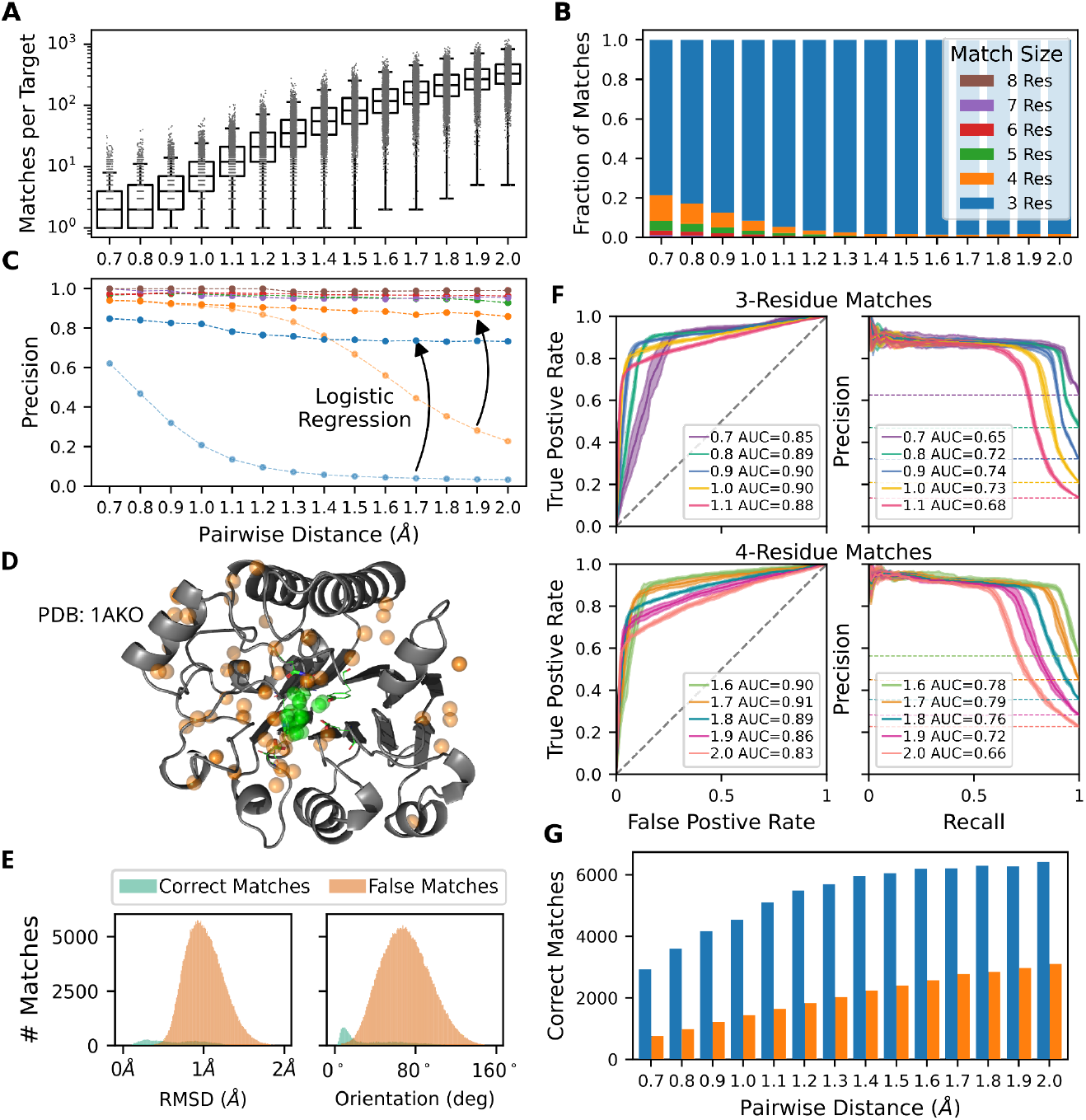
(A) The distribution of the number of matches per target on a logarithmic scale and (B) their relative residue composition broken down by pairwise atom distance (Å). (C) Precision of template matching by pairwise atom distance. For 3- and 4-residue matches (blue and orange lines, respectively), logistic regression models filter poor matches and improve precision. Colours in panels B, C and G indicate the number of residues in a match. (D) Example of correct (green) and false (orange) matches at 1.5 Å against 1AKO structure. Spheres indicate the centre of mass of a match. (E) Distribution of RMSD and residue orientation values for correct (green) and false (orange) matches at 1.5 Å. (F) ROC and precision-recall curves for a 5-fold cross validation of logistic regression models trained at each pairwise distance to predict correct matches based on RMSD and residue orientation for 3- and 4-residue matches. Bold lines show the mean curve over all models with shaded areas indicating the standard deviation. Mean AUC values are shown. (G) Bars indicate the total number of correct matches with 3- or 4-residues after filtering by logistic regression models.

We computed precision (fraction of correct matches) for varying template sizes against the pairwise atom distance (Fig. 2C). Templates with five or more residues were nearly always correct at all pairwise distances, with few exceptions attributable to incomplete annotations. Thus, larger templates were shown to maintain their selectivity likely because the constraints of specified residue and atom types present in close proximity is sufficient. Similarly, the correct fraction of 4-residue matches was much higher than that of 3-residue matches, though precision declined with higher pairwise distances. From these data we estimate the likelihood of 5+ residue matches to occur by chance to be almost zero. We therefore focused on distinguishing correct, 3- and 4-residue matches from false ones. For example, 29 correct (green) and 61 false (orange) matches to the structure of the *E. coli* exonuclease III (PDB: 1AKO*(31)*) were found (Fig. 2D). While correct matches clustered together in the active site, false matches were distributed over the entire structure.

True recall is difficult to estimate, as without entries in the M-CSA, it is very difficult to define instances of catalytic similarity. Hence, performance evaluation was restricted to assessing precision (i.e. the proportion of matches correctly detecting the true catalytic site). Considering the heterogeneous template selectivity, our evaluations are intended to apply across templates.

### Developing match quality metrics

To reduce the number of false positive matches, we developed geometric match quality metrics. In the original Jess publication, RMSD calculated by superposition of all target and template catalytic atoms was identified as a good measure of match quality when accounting for size of the template. Since intermolecular interactions with the exception of coulomb forces are anisotropic (i.e. depend on relative angles), we hypothesized that comparing the orientation of residues between the target and the template would complement RMSD. To test this, we calculated orientation vectors for each residue which we then compared by calculating the angle between corresponding template and target residues. The arithmetic mean of pairwise inter-template-target angles was chosen as a measure of residue orientation difference (Fig. 2E showing matches at 1.5 Å). As expected, we found that for correct matches, both measures correlated well at all pairwise distances with Pearson’s r coefficients ∼0.8. By contrast, the higher the pairwise distance, the lower the correlation was for false matches, indicating that RMSD and residue orientation capture different features as structural differences between template and target increase (see Fig. S6–S8 for distributions and correlations of 3-residue data at 0.9 Å and 4-residue data at 1.7 Å).

### Distinguishing correct and false matches

To systematically distinguish correct from false matches based on these properties, we trained logistic regression models on RMSD and residue orientation of 3- and 4-residue matches separately at pairwise distances between 0.7 Å and 2.0 Å. Match data was split for a five-fold cross-validation after randomly shuffling into train and test sets stratified to contain similar proportions of correct and false matches. Performance of all 5 models from cross-validation was evaluated using ROC and precision-recall curves (Fig. 2F, for clarity, only selected pairwise distances with the best performances are shown). The decision threshold for identifying correct predictions was set to maximise the Matthews correlation coefficient (MCC, Fig. S9). For Enzyme Motif Miner, at a given pairwise distance, a prediction is made by majority vote of all 5 models. As expected, we found that there is a tradeoff between precision and the total number of predicted correct matches (Fig. 2G). Based on our findings, we recommend a pairwise distance threshold of 0.9 Å for 3-residue matches and 1.7 Å for 4-residue matches, while for 5+ residue templates 2.0 Å, maximising the number of matches is most suitable.

Hydration, pocket volume and solvent accessibility also play an important role in catalysis. We investigated the predictive power of solvent accessibility as an additional feature but found that this only offered a minor improvement which did not compensate for the additional computational expense.

With these improvements in precision, Enzyme Motif Miner can, in some cases, identify additional catalytic residues in proximity to the annotated catalytic site as well as some metal binding sites - a feature which is useful in the context of predicted protein structures commonly lacking cofactors. In the supplement we discuss examples of both, illustrated by means of a bacterial zinc beta-lactamase and a bacterial alcohol dehydrogenase.

### Performance compared to sequence and structure identity

We next assessed how Enzyme Motif Miner compared to sequence and structure-based homology searches. For the latter, we took inspiration from ‘Foldseek’, a popular method for structure alignment, which uses a variational autoencoder to predict a 20-letter alphabet sequence along the protein chain called 3Di sequence*(14)*. This 3Di sequence represents the protein structure based on the geometry of closest non-sequential neighbouring residues and enables fast k-mer based alignment algorithms. We compared Enzyme Motif Miner to pairwise global amino acid and 3Di alignment (see the *Validation dataset* section of the methods for details) on our validation set of highly divergent enzymes at various maximum global sequence identity thresholds to predict correct or false matches based on their sequence identity. For this purpose, datasets at 0.9 Å for 3-residue matches and 1.7 Å for 4 residue matches were selected since these are closest to a dataset with similar proportions of positives and negatives.

We found correlations between sequence identity and RMSD for correct matches (Pearson’s r: -0.23 and -0.37 for amino-acid and 3Di global identity respectively) (Fig. S10). We observed that models trained on RMSD and residue orientation far outperformed sequence identity models. Below about 27.5% amino-acid sequence identity and 45% 3Di identity, sequence identity models performed no better than random (Fig. S11 for ROC curves). These data show that for evolutionarily distant enzymes, Enzyme Motif Miner remains predictive by leveraging catalytic motifs which are strongly conserved, instead of sequence information.

### Identifying function and convergence or distant divergence

For targets and templates annotated with complete EC or CATH numbers, we compared their similarity by agreement in the respective hierarchical levels of these classification schemes. EC encodes enzyme reaction chemistry and CATH describes protein domain structure (Fig. 3). Correct matches with a high CATH similarity (orange and blue fractions) likely indicate homology, while false matches with high similarity are random matches in non-catalytic parts of a homologous structure. Among the latter, CATH matches at only the 1st or 2nd level are between proteins which are unrelated or evolutionarily extremely distant. We further focused on the small fractions of correct matches assigned into different CATH superfamilies, as these intriguingly represented proteins with similar motifs of either convergent or divergent origin. In any case, the existence of such matches further corroborated that Enzyme Motif Miner is sensitive to similarities not detectable in sequence and overall structure, but only in their catalytic site geometries.

**Figure 3.**
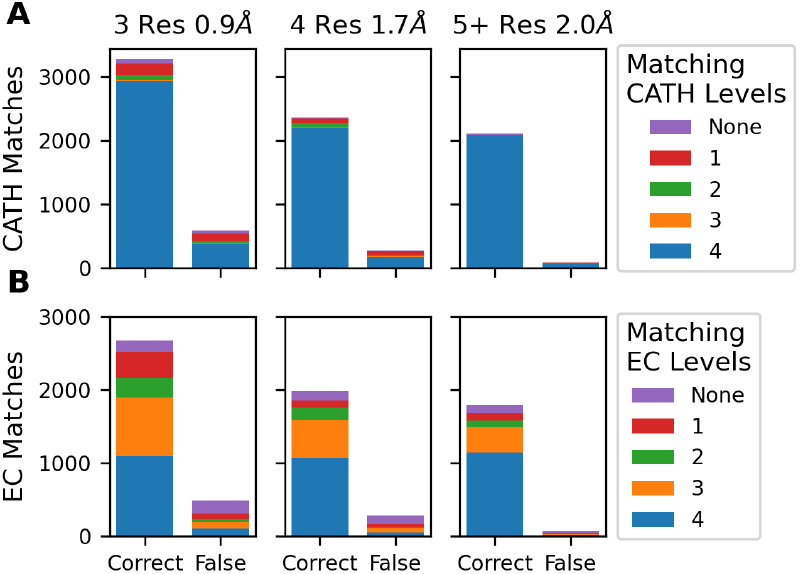
Predicted matches split by correct and false matches and coloured by their level of CATH (A) or EC (B) similarity. Only matches where both target and template had complete EC numbers are considered. Data is shown for 3-residue matches at a pairwise atom distance of 0.9 Å, 4-residue matches at 1.7 Å and larger matches collected at 2.0 Å.

A majority of correct matches shared a 3rd (orange) or 4th level (blue) EC similarity (Fig. 3B). This is in agreement with the CATH similarity distributions (Fig. 3A), highlighting the connection between conserved catalytic sites and enzymatic mechanisms. Generally, lower similarities to EC annotations as compared to CATH annotations were observed, which can be explained by slight changes in substrate binding pockets leading to different enzymatic function even though the mechanisms and relevant catalytic residues stay largely the same. This is particularly the case for the 4th EC level which is often substrate or product specific. This is in line with our expectations for smaller templates representing partial catalytic sites which may facilitate similar steps in different catalytic mechanisms such as proton transfer or metal-coordination. Such stepwise divergence is not necessarily well reflected by EC classification. For these reasons smaller templates are likely suitable for detecting very distantly divergent homologs or even examples of structural convergence following functional convergence. By selecting matches with low CATH similarity but high EC similarity and comparing the sequence order of matching residues, we aim to distinguish convergent and divergent evolution. We further explored some examples of possible convergence (Fig. 4,5).

**Figure 4.**
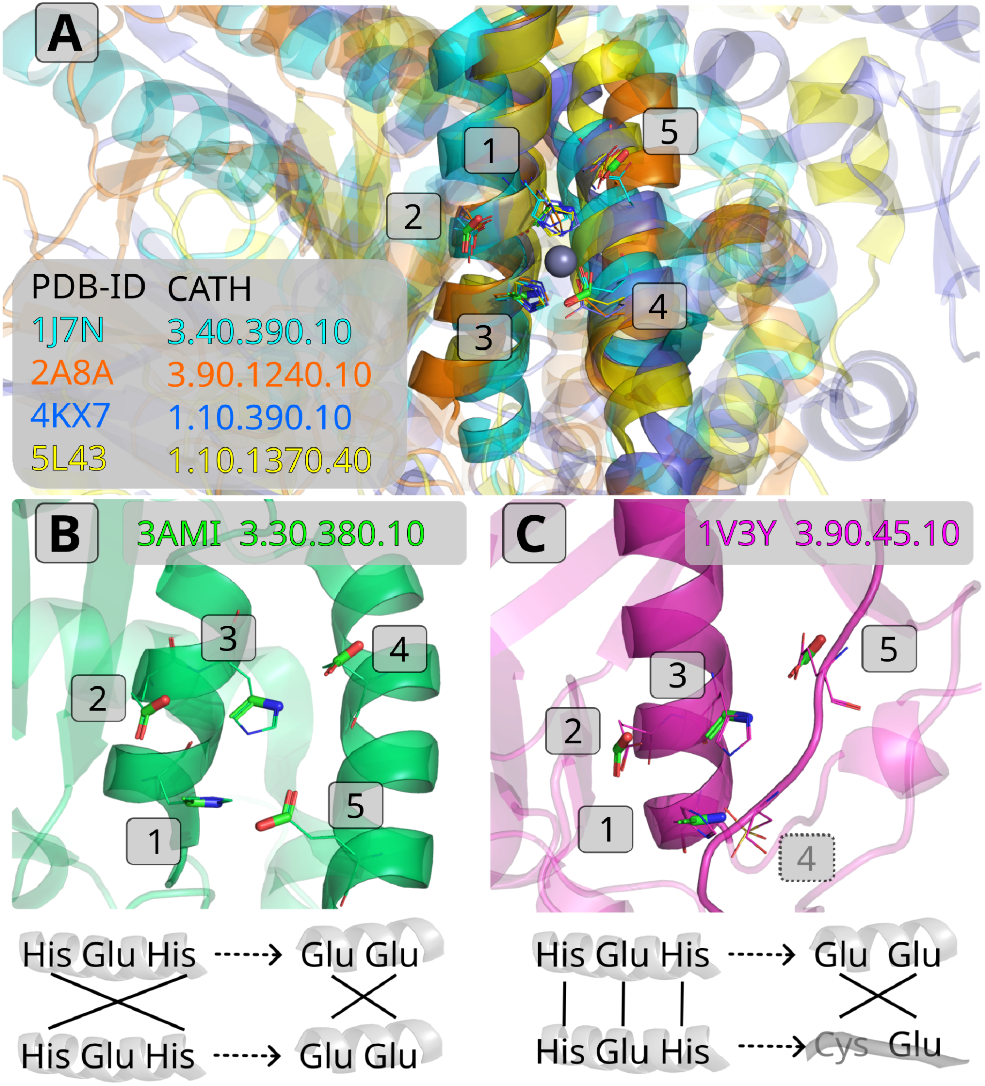
Enzymes with different CATH domains were identified using 4- and 5-residue templates from (green sticks, PDB: 3AMI(32), M-CSA 657). Each CATH domain is coloured differently. Numbers 1-5 denote the relative sequence order of matched residues in each structure. All panels show the active site arrangement in the same orientation. (A) PDB structures 1J7N, 2A8A, 4KX7 and 5L43 are aligned on matching catalytic residues (lines), revealing two anti-parallel helices. All structures in panel A shared the same relative sequence order of catalytic residues. (B) The template structure 3AMI shared the same catalytic arrangement between two homologous helices, however both helices were inverted (bottom) with respect to the consensus sequence (top) of structures in panel A. (C) The PDB structure 1V3Y matched four of the catalytic residues. For 3AMI and 1V3Y the change in sequence order is visualised below. Lines indicate corresponding residues which are shadowed by their respective secondary structure.

**Figure 5.**
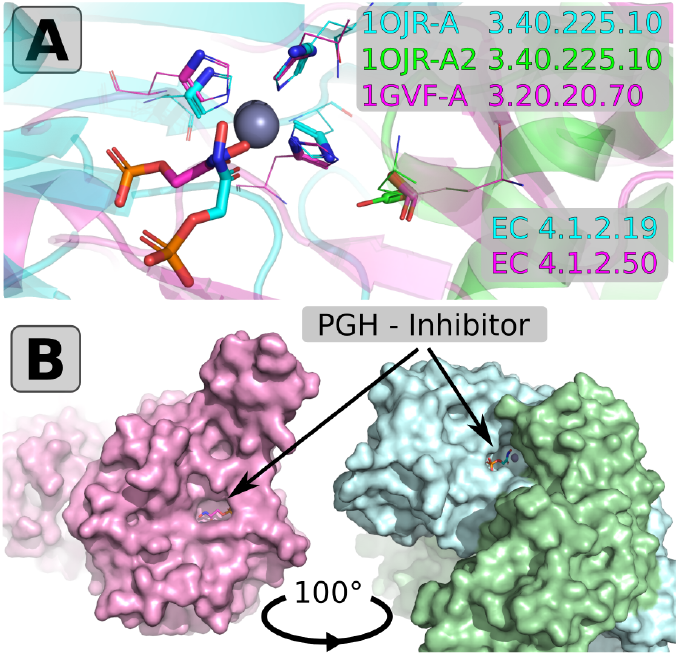
(A) A view of the active sites of the template 1OJR and 1GV3 after aligning matched residues (sticks).The catalytic residues are shown as lines. 1OJR is a homo-tetramer with the active site split across two chains (Chain A green, Chain B cyan). The same arrangement of residues was found in the single chain active site of 1GVF (magenta). CATH numbers are indicated. Both structures share the EC number 4.1.2.- and are bound by the inhibitor phospho-glycolohydoxamic acid (PGH - shown as sticks) in similar positions. 1G75(38) was superimposed onto 1OJR to include the inhibitor (global RMSD 0.31 Å). (B) A surface representation of both proteins but with views rotated by ∼100° show that active sites are accessible from opposite directions.

One case involved 27 matches identified using three- to five-residue templates derived from the *Sphingomonas sp*. M16-B metallopeptidase structure (PDB 3AMI, Fig. 4). Its active site comprises two histidines and three glutamates. His1, His3, and Glu4 (or Glu5 in 3AMI) coordinate zinc, while Glu2 acts as a nucleophile to activate a water molecule for peptide bond hydrolysis, and Glu4 (Glu5) stabilizes His1*(32,33)*. These 27 matches span seven distinct CATH superfamilies, all dissimilar from 3AMI. Among these, we found examples (1J7N, 2A8A, 4KX7, 5L43) with different domain architectures, all aligned with the 3AMI template residues (Fig. 4A,B). Functionally, all are metallo-peptidases (EC 3.4.24.- or 3.4.11.-). Aligning catalytic residues revealed a conserved secondary structure: two antiparallel α-helices around the catalytic site, with His1, Glu2, His3 on one helix and Glu4/5 on the other. Apart from these helices, no structural conservation was found. Even the helices themselves show major variations such as an 81-residue insertion in 2A8A between Glu4 and Glu5. Notably, none of the matches preserve 3AMI’s residue order; the sequence in each helix often appears inverted, making descent from a common ancestral sequence unlikely.

Another intriguing match detected with this 4-residue template was the peptide deformylase 1V3Y*(34)* from *Thermus thermophilus* with the EC number 3.5.1.88 (Fig. 4C). This enzyme uses a cysteine in place of glutamic acid as a third metal coordinating residue. While the enzyme cleaves an N-formyl bond, the mechanism is otherwise similar to other metallopeptidases. The sequence order of the remaining left helix mirrored that of 3AMI but was swapped for the other two residues.

An interesting example of potential convergence is the four-residue template from *E. coli* L-Rhamnulose-1-phosphate aldolase (1OJR*(35)*, M-CSA 645) matched to tagatose-1,6-bisphosphate aldolase (1GVF*(36)*) (Fig. 5A). Although the active site residues are correct, orientation differences led the match to be filtered out. 1OJR’s active site spans two chains (green and cyan), whereas 1GVF uses one. Both share EC levels 4.1.2.-, coordinating three His residues with Zn^2+^ and a Glu acting as Brønsted acid during opening of the sugar ring. Functional convergence seems likely since both enzymes are inhibited by phospho-glycolohydoxamic acid despite differing surrounding structure, while aligned active sites face opposite directions (Fig. 5B).

These examples demonstrate the utility of catalytic site matching to reveal local structural and functional relationships that are invisible to sequence or domain classification, uncovering both distant divergence and convergent mechanisms. For further examples, we have compiled tables of all correct matches among PDB structures with different CATH domain folds that are either divergent or possibly convergent (Zenodo*(37)*).

### Applications to predicted protein structures

We evaluated Enzyme Motif Miner on Alphafold2*(2)* predicted protein structures of the Swiss-Prot*(25)* human proteome for its ability to distinguish enzymes from non-enzymes (assessed by comparison to Swiss-Prot-provided enzyme annotations). Of the 20.406 human proteins (Dec. 20 2024), a quarter (5531) are annotated as enzymes according to Swiss-Prot. Overall, 75% of total matches by Enzyme Motif Miner were to structures also annotated as enzymes. Enzyme Motif Miner yielded matches to 2335 or 42.0% of all annotated enzymes while matching only 6.4% of non-enzymes. For 1431 enzymes, we were able to validate Enzyme Motif Miner matches with active site annotations in UniProt*(39)*. Out of all the human enzymes with active site annotations Enzyme Motif Miner detected, 83.7% were matched on residues belonging to the annotated catalytic site, demonstrating that precision is maintained for predicted protein structures.

To follow up on our matches to human enzymes, we compared the agreement in EC number classification between the template and the enzyme. We find that for nearly 60% of 3- and 4-residue matches, the EC number is correct to the 3rd level while for 5+ residue matches, this fraction approaches 80% (Fig. 6) With our template library we still missed matches to catalytic sites in the remaining 3220 human enzyme structures in Swiss-Prot. Based on comparisons of EC and Pfam annotations as well as homology to the M-CSA we might have plausibly expected matches to about a quarter of these structures (Fig. S12 for a breakdown). These false-negative predictions may be due to AlphaFold2 predicted structure quality being too low, conformationally dissimilar or with an active site only found at the interface of multiple chains. Further matches may have been found for another third of missed enzymes if coverage of the M-CSA would be extended to larger parts of the PDB. For the remaining 35%, there is a lack of experimental evidence (location of the active site, function and structure) precluding template generation. We further investigated why some non-enzymes were matched with the result that many of these matches were mostly composed of metal coordinating residues which also feature in many catalytic sites (Fig. S13).

**Figure 6.**
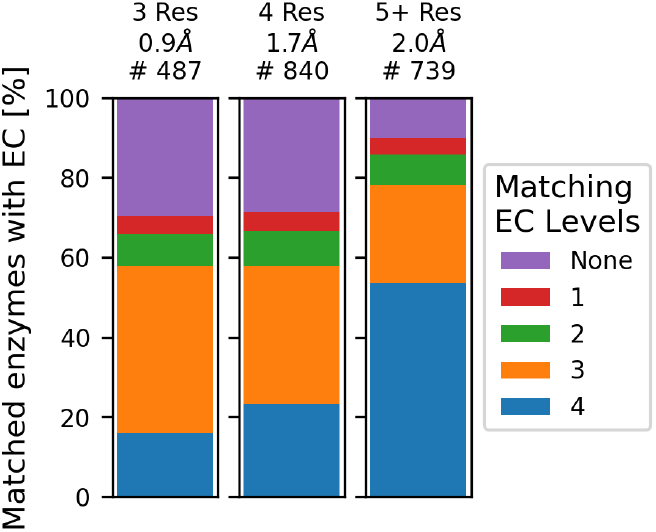
On a set of Alphafold2 predicted structures of human enzymes for which Swiss-Prot reviewed annotations were available as a ground truth, matches were found to 2068 enzymes with at least 3rd level EC annotations. This figure illustrates the relative fractions of matches in which the EC number between the human enzyme and matched M-CSA template overlap at a given EC level. Matches were found at the previously determined pairwise distances (after filtering by logistic regression models).

To test template matching performance on the fringes of structural similarity we turned to a set of 7427 novel predicted protein folds which fall outside the CATH hierarchy identified in the encyclopaedia of domains*(40)*. Despite expecting no homology to any PDB structures, we were still able to find 766 matches to 442 of these novel domains. Most of these matches can be attributed to metal binding sites. We chose to specifically investigate 27 domains with 5+ residue matches. Among those, we identified one putative metallophosphoesterase which aligned on seven metal coordinating residues. Remarkably we found significant similarity around the active site despite considerable divergence outside the active site. A further cluster with similarity to phospholipase C templates was identified. Additionally, we found eight domains with similarity to metalloproteases featuring two antiparallel helices (Fig. 4 for reference). Only four domains were identified by 5+ residue matches to non-metal binding sites (Fig S14). With RMSDs of 1.2-1.7 Å none of them showed particularly good alignment. Nonetheless, three of these matches constituted plausible binding pockets which based on visual exploration might have catalytic potential (Fig. S15).

## Conclusions

With Enyzme Motif Miner we have developed a robust and very fast method to geometrically match known catalytic templates in protein structures. Like many previous tools, this approach initially suffered from false-positive matches occurring by chance to functionally dissimilar protein sites. While larger, more constrained templates are more selective, this problem is particularly acute for smaller matches to 3- and 4-residue templates. We addressed this issue of spurious matches by filtering with logistic regression classifiers based on RMSD of match superposition and residue orientation angles. This filtering does however come at the cost of reducing sensitivity.

We also demonstrated that Enzyme Motif Miner is a useful tool for detecting structural convergence of catalytic features in otherwise non-related enzymes. Nonetheless, it remains challenging to distinguish between similarity at the functional level and the catalytic or mechanistic level, furthermore complicating prediction of functional enzyme properties from template matches alone. Enzymes may well share functional 3D residue arrangements or individual catalytic steps even if their overall reactions and thus substrate and product are dissimilar. Substrate specificity is often guided more by binding residues in the first and second shell rather than by active site residues directly involved in catalysis. This phenomenon is for example observed for metal coordinating residues, which are distinct from other catalytic roles. For these reasons, Enzyme Motif Miner is not intended to predict overall reactions or substrate profiles.

Insights from Enzyme Motif Miner can be fully traced back to their original data sources. Our template-matching approach is also highly efficient, outpacing recent AI-based function prediction tools such as DeepFri*(41)* and PARSE*(42)* by two orders of magnitude. Moreover, both PyJess and Enzyme Motif Miner provide seamless interoperability through a modern Python API. As the structural proteome is expanding extremely rapidly due to AI structure prediction algorithms, Enzyme Motif Miner is built to scale alongside the development of such resources. To showcase its utility, we validated Enzyme Motif Miner on datasets of predicted protein structures and demonstrated retrieval of enzymes and their EC classification in the human proteome as well as its applicability to the exploration of entirely novel folds on a dataset from the encyclopaedia of domains.

An important limitation of this work is its reliance on annotations from the M-CSA for both generating the template library as well for classifying matches as correct and false during validation. Using a fixed library of templates curated from literature sources, our approach cannot be used to extrapolate beyond the well known and preferably experimentally characterised enzymes to make novel associations. Coverage of catalytic space in the M-CSA is likely far from comprehensive even though most currently described reaction mechanisms up to the 3rd level are covered. Therefore, novel catalytic structures or related but modified catalytic structures or larger conformational deviations without close resemblance will not be detected. We still lack matches for slightly over half of human enzymes in predicted protein structures. While we attribute this primarily to a lack of template coverage, protein prediction quality especially with regard to side chain orientation of catalytic residues may play a role as well.

The underlying principles of this work can also be applied to other types of functional protein sites. Recent work has leveraged template matching to detect intramolecular isopeptide bonds*(43)*. As drug or ligand binding sites encompass much more and often unknown conformational variation this would not be a straightforward task however. In the case of protein-protein interactions, approaches encoding signs of coevolution in multiple sequence alignments, would likely yield much better results.

The most obvious applications of Enzyme Motif Miner lie in improving the knowledge driven catalytic annotation of uncharacterised protein structures and surveying similarities between active sites. Besides annotation, we showcased how to gain insights into the relatedness of enzyme catalytic sites. Mapping the similarity of catalytic residues in combination with sequence and fold homology could improve our understanding of how enzymes diversify in terms of catalytic function. For example, through comparisons to CATH annotations we found many cases of divergence surrounding metal-dependent protein hydrolases catalysing a reaction centred on two antiparallel helices.

Using catalytic templates is one approach to understanding catalysis and enzyme function and evolution. Given that available protein sequences currently vastly outnumber available structures, this sequence information should also be leveraged, for example, to investigate the evolution of the enzyme. Detecting enzyme relationships by template matching is made more complex by the variety of side chain conformations and flexibility seen in some active sites. Such flexibility does not affect sequence analysis. Therefore, we recommend that template analysis, whilst more detailed and able to locate the active part of the enzyme, is always complemented by sequence-based analysis.

## Methods

### Validation dataset

We aim to create a set of non-redundant protein structures that are homologous to one of the reference entries in the M-CSA, to explore if Enzyme Motif Miner can detect these relatives. PDB enzyme structures with annotated catalytic residues were obtained from the Mechanism and Catalytic Site Atlas (M-CSA)*(4)*. Sequence homologs at the chain level based on the representative PDB structure of each entry to other structures in the PDB were identified using HMMER*(44)*. Sequences of structures with an E-value lower than 1x10-6 were classified as homologs. Catalytic residue annotations were transferred across homologs based on multiple sequence alignments. Across all entries in the M-CSA 53,047 homologs with 113,055 chains were annotated with at least three catalytic residues. For validation, only chains from PDB entries active as of June 2024 with resolution better than 3 Å and R-factor better than 0.25 with sequence lengths between 40-10,000 residues were selected using PISCES*(45)* annotations. For NMR structures containing multiple models, only the first model was included. The sequences of these chains in the PDB were hierarchically clustered using CD-hit*(46,47)* in increments of 10% sequence identity down to approximately 40% maximum sequence identity with word sizes from 5 down to 2. This resulted in a total of 3751 different target structures. Pairwise global sequence alignments were calculated between all target chains using the Needleman Wunsch algorithm*(48)* (gap open: 11, gap extend 1) as implemented in PyOpal (https://github.com/althonos/pyopal). The BLOSUM62 matrix was used for amino-acid sequences and the ‘foldseek’ substitution matrix for 3Di sequences (*https://github.com/steineggerlab/foldseek/blob/master/data/mat3di.out)(14)*. 3Di sequences encoding protein structure were calculated through a Python implementation of the ‘foldseek’ VAE (https://github.com/althonos/mini3di).

Modified or unnatural amino acids not included in the BLOSUM62 matrix were treated as unidentified residues. Structures may contain residues mutated with respect to their sequence record in UniProt but catalytic annotations are based on the sequence of the structure record in the PDB. Annotations for EC, CATH and UniProt*(39)* were taken from SIFTS*(49,50)*.

### Predicted protein structure datasets

We collected a list of 20,406 Swiss-Prot*(25)* reviewed proteins of the human proteome from UniProt*(39)* and downloaded all available structures from the AlphafoldDB*(2)*. Since proteins larger than 2400 residues are chunked into segments of 1400 overlapping by 200 residues, we retrieved 23,116 structures. Template matching was performed using default settings matching 3-residue templates at a pairwise atom distance of 0.9 Å, 4-residue matches at 1.7 Å and larger ones at 2.0 Å. Smaller matches are skipped if larger matches could be identified. We excluded residues with a pLDDT below 70 from being matched. Enzymes were identified by their GO annotations (term: GO:0003824) and, if available, active site annotations in UniProt. For analysing novel folds in the encyclopaedia of domains, we downloaded all 7,427 novel structures*(40)*. Identical parameters to above were applied for identifying matches. Data for both analyses is available on Zenodo*(37)*

### Templates

A template contains a set of catalytic residues and their 3D coordinates while specifying a set of constraints such as interchangeable amino acid types and allowable structural flexibility. Templates in our library contain up to 8 residues. Each residue is represented by three functional atoms (Fig. S1), according to its function as annotated in M-CSA. Residues which interact through both side- and main-chain atoms are represented by six functional atoms. For 813 of the current 1004 entries (81%) in the M-CSA, templates have been derived using our previously published CSA-3D package*(6)*. Here, homologous PDB structures were clustered and the representative members were collected as templates, each describing a consensus active site conformation. Thus templates account for known differences in conformation. A given template residue may specify a small selection of chemically equivalent amino acids (e.g. Asp-Glu, Ser-Thr-Tyr) if such substitutions are observed in homologous enzymes. This way a template’s constraints may account for both divergence through conservative missense mutations as well as functional convergence. Larger templates are themselves subdivided into smaller composite patterns of fewer residues describing partial active sites, identified by applying a k-means algorithm in 3D. The exact methodology is described by Riziotis et al.*(6,29)*.

Thus by subdivision of larger templates and considering alternate catalytic conformations, a total number of 6780 templates from 1412 PDB structures across 762 M-CSA enzyme families were used for analysis. Only the number of unique and defined residues interacting through their side chain in a template are counted towards its size. Thus, residues with six functional atoms are counted only once and residues allowed to match any amino acid type are not counted. This was done in order to make selectivity more comparable to template size as atoms allowed to match to backbone atoms of any residue type were observed to be much less selective. The size distribution of our template library as given by unique, specific residues is shown (Fig. S2). While we provide our library of templates, users may also use their own templates. Templates make use of a modified PDB-like format.

Template annotations such as EC number, CATH accession and InterPro*(40*) annotations were collected from the M-CSA. The residue order and the adjusted number of unique, specific residues in each template are calculated alongside the orientation of each template residue given by an amino acid type dependent vector. To analyse EC coverage, a record of EC numbers was collected from KEGG*(30)* (https://www.genome.jp/kegg/annotation/br01800.html).

### PyJess

We developed PyJess (https://github.com/althonos/pyjess) as a stand-alone package to provide Cython bindings and a Python API for Jess in a thread-safe manner allowing for parallelized matching. Further, we made several algorithmic optimisations to PyJess as outlined below. For improved interoperability, PyJess handles both PDB as well as CIF file formats (using GEMMI*(52)*) and can load molecules from other commonly used Python libraries. PyJess translates a template into a set of constraints for maximum pairwise atom distances, residue and atom type which together represent a 3-dimensional kd-tree. By iteratively expanding any partially matching solution until all constraints are satisfied, the PyJess algorithm can efficiently query a large number of structures for similarity to the template. Searching for a template in the structure from which it was derived will always return the template atoms except for rare cases of residues chemically modified for crystallisation. Since the algorithm only compares pairwise distances and atom/residue identifiers, no computationally expensive alignment is required. Only after a solution satisfying all template constraints is found, is the match transformed into the coordinate reference frame of the template and the two structures are superposed.

In addition to a library of targets and templates, three parameters controlling the search are given to PyJess. PyJess is run using specified cutoffs for RMSD, pairwise distance between atoms and B-factor. Only matches with an RMSD lower than the RMSD cutoff (defined below) in ångström (Å) are reported. As such, this threshold does not affect the search algorithm in any way. Pairwise distance between atoms controls the internal search radius around each atom within which atoms are allowed to match. Finally, we implemented a B-factor cutoff in order to apply per-residue cutoffs like pLDDT scores in predicted protein structures, where they are stored in the B-factor column. Atoms match only if the B-factor of the atom in the protein structure is above or better than this threshold.

### Performance optimizations

PyJess introduces a series of optimizations over the previous Jess algorithm, ensuring consistent results while improving the efficiency of structural template matching. A major improvement was made to construction of the k-d tree, which is used to partition a structure into regions for faster atom retrieval. PyJess now reorders atoms using QuickSelect in linear time, substantially reducing the cost of building spatial partitions of the k-d tree for large structures. Query performance is further improved by an approximate annulus-intersection test. Rather than immediately computing exact euclidean distances between bounding boxes and annuli, PyJess performs a coarse check using bounding-box overlaps. Only nodes that pass this filter undergo the more expensive geometric evaluation, eliminating most unnecessary distance calculations. Backtracking in the k-d tree is also reduced by selecting an optimal matching order such that template atoms are sorted by the size of their candidate sets so that the most restrictive atoms are matched first, which reduces the number of failed partial matches and the associated k-d tree lookups by a factor of 10.

We also introduce indexing a query molecule by residue names, which benefits a majority of templates in our library. Additional optimizations include inlining and concretizing routines, fixing the implementation to three dimensions, recycling scanner memory, and replacing generic case-insensitive string comparisons with a specialized routine. A full description of implemented optimizations is provided at https://pyjess.readthedocs.io/en/latest/guide/optimizations.html.

### Enzyme Motif Miner

With Enzyme Motif Miner we provide both an internal Python API and an easy to use command line interface wrapping PyJess. A library of catalytic templates generated from the M-CSA is supplied. Templates are matched in descending size order which allows the user to easily run different parameters depending on the template size. Secondly, should a larger template match the target structure, searches against smaller templates can be skipped to speed up the search process. For validation, this option was disabled to compare template performance based on size. From this validation process, logistic regression models were fitted to distinguish correct and false matches at each pairwise atom distance for templates composed of 3- and 4-residues.

Predictive models at pairwise atom distances of 0.7 Å to 2.0 Å in increments of 0.1 Å were fitted. Larger pairwise distances become increasingly computationally expensive and unreliable, while stricter cutoffs do not require much validation. By default, as well as during validation, a B-factor cutoff was not applied since only structures with resolutions better than 3 Å and R-factor better than 0.25 were selected. Based on prior experiments we chose a generous RMSD reporting threshold of 2.0 Å. For each target structure only the best match with the lowest RMSD value to each template was reported. Metadata about matches is saved in tsv files while template-target superpositions for all matches to a target are stored in PDB file format. Data from all validations is available on Zenodo*(37)*

### Data analysis

The aim was to assess whether Enzyme Motif Miner could correctly identify catalytic sites in distant template homologues and the impact of changing pairwise distance cutoffs on their detection. As PDB structures obtained through the M-CSA often contain multiple chains, only matches including the non-redundant chains selected for analysis were considered but matches across chain interfaces were still allowed as long as at least a part of the match was to the selected set of target chains. We defined a template to have correctly matched the active site location if at least three of the matched residues are annotated as catalytic. Note that for some proteins in our target dataset catalytic annotations may be incomplete.

In case of EC and CATH numbers, annotations for target structures were compared to template annotations. Results were analysed using Python 3.13.1. Data handling and visualisation was done using Numpy v2.1.3*(53)*, Scipy v1.15.1*(54)*, Sklearn v1.7.0*(55)*, Matplotlib v3.10.0*(56)*, Plotly v6.2.0*(57)*, Polars 1.29.0*(58)*, Pandas v2.2.3*(59)* and Seaborn v0.13.2*(60)*. Performance was assessed using Valgrind v3.25.1*(61)* and KCachegrind v23.08.5*(62)*. Final performance tests were conducted with hyperfine v1.19.0*(63)*. 3D-molecular figures were generated using an open source version of Pymol obtained from https://github.com/schrodinger/pymol-open-source/.

### Calculating match orientation

For each 3-atom residue in the template, a vector in 3D space calculated according to the amino acid type was defined. This vector approximates the spatial orientation of the functional group of the residue. For residues in which two of the functional atoms are indistinguishable by element such as for arginine, glutamic acid, aspartic acid and phenylalanine, the vector was drawn from the terminal side chain carbon to the midpoint between the two identical elements (Fig. S1 for atom pairs which define the orientation vector for each amino acid). The analogous vector for the corresponding matched residue in the target was then likewise calculated. Principally, angles can then be derived from these vectors by comparing corresponding vectors between template and target or by comparing all pairwise angles within each structure. After transforming the target coordinates into the template reference, vectors within a match can be compared. Such inter-template-target angles can then be combined into a simple arithmetic mean. Other means such as the RMS seemed to yield similar results. For this reason the arithmetic mean, which is the most straightforward approach, was preferred. Residue orientation refers to the arithmetic mean of inter template-target angles between residues.

## Acknowledgements

We would like to thank both the Thornton and Zeller groups for many stimulating discussions. In particular we are very grateful to the entire M-CSA team for maintaining, developing and curating the Mechanism and Catalytic Site Atlas. We thank Jonathan Barker for his clever engineering in developing Jess as maintainable open source software. We would also like to express our gratitude to Neera Borkakoti and Roman Laskowski for their support and the many incredibly helpful discussions which have shaped this project.

## Author Contributions

Conceptualization: JMT

Data Curation, Formal Analysis, Visualization: REH

Software: REH, ML, AJMR

Supervision: IGR, GZ, JMT

Methodology: IGR, GZ

Resources: AJMR

Funding: REH, GZ, JMT

Writing - Original Draft: REH

Writing - Review & Editing: REH, IGR, ML, AJMR, GZ, JMT

## Data and materials availability

Protein Structure Data is available from the PDBe. Information on catalytic sites was derived from the Mechanism and Catalytic Site Atlas and is available for download at https://www.ebi.ac.uk/thornton-srv/m-csa/download/.

PyJess is available at https://github.com/althonos/pyjess and can be installed via PyPi from https://pypi.org/project/pyjess/.

Enzyme Motif Miner is available at https://github.com/rayhackett/enzymm and can be installed via PyPi from https://pypi.org/project/enzymm/.

The original Jess can be accessed at https://github.com/iriziotis/jess/ All code is available under MIT licence.

Results from validations are available through Zenodo*(37)*

Supplementary Information

