## Supplementary Information for "Investigating Enzyme Function by Geometric Matching of Catalytic Motifs"

### Supplement

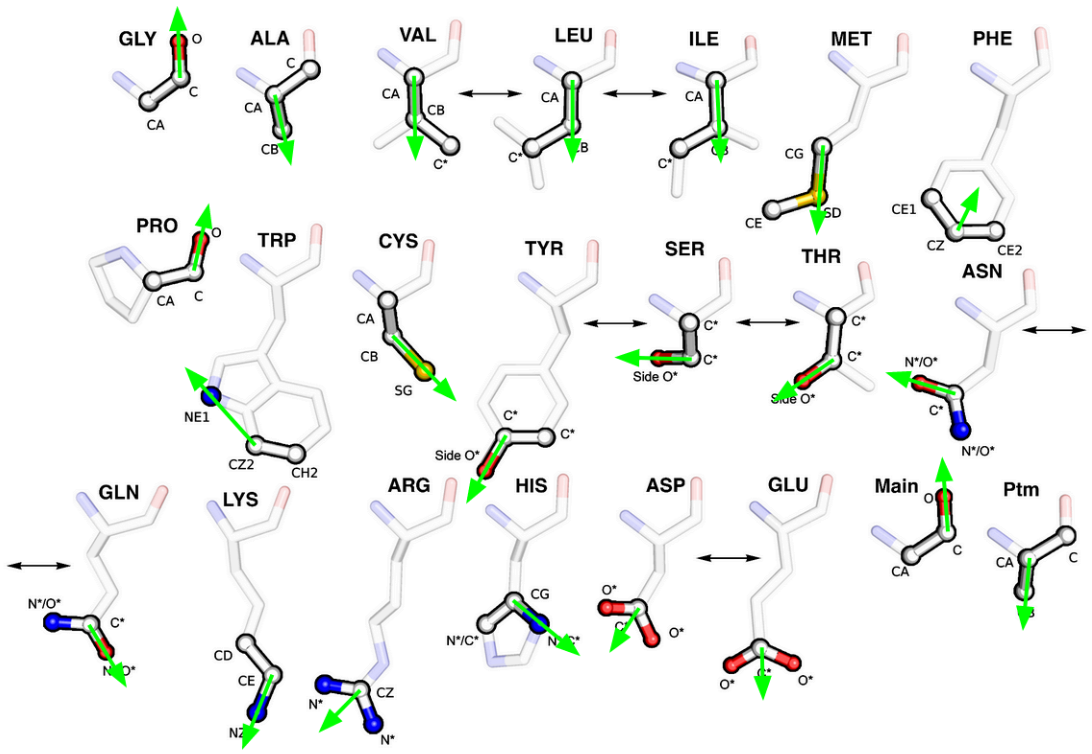

GLY: (C, O), ALA: (CA, CB), VAL: (CA, CB), LEU: (CA, CB), ILE: (CA, CB), MET: (CG, SD), PHE: (CZ, mid), TYR: (CZ, OH), TRP: (CZ2, NE1), CYS: (CB, SG), PRO: (C, O), SER: (CB, OG), THR: (CB, OG1), ASN: (CG, OD1), GLN: (CD, OE1), LYS: (CE, NZ), ARG: (CZ, mid), HIS: (CG, ND1), ASP: (CG, mid), GLU: (CD, mid), PTM: (CA, CB), ANY: (C, O)

*Figure 1. Each residue in a template is represented by three functional atoms each. Atoms which define a residue are emphasised. This figure was adapted from Riziotis et al.(1). For each 3-atom residue in the template a vector depending on the amino acid type is defined. Green arrows indicate this residue orientation vector. Atom names as defined by the PDB are shown. Mid refers to the euclidean midpoint between the two other atoms.*

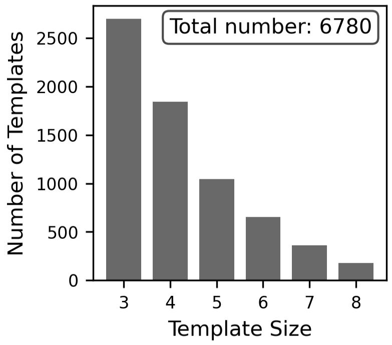


*Figure 2. Templates range in size from three to eight residues. The size distribution of the template library is shown as a bar chart. Only unique residues matching a defined set of amino acids are counted towards the size of a template.*

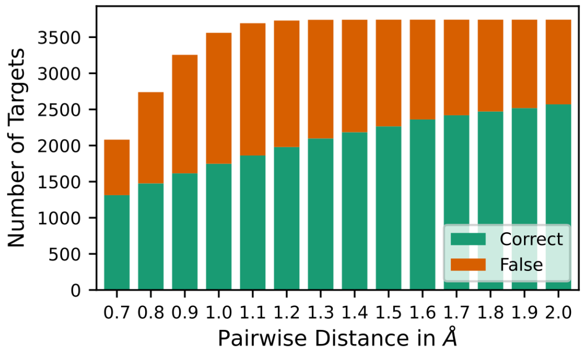


*Figure 3. In total 3751 target structures were searched. At pairwise distances above 1.2 matches were found for nearly every target structure. However, after distinguishing between correct and false matches, only for around ⅔ of the structures, was at least one correct match identified. This bar plot shows the number of targets for which any match was found, split by whether at least one match was correct.*

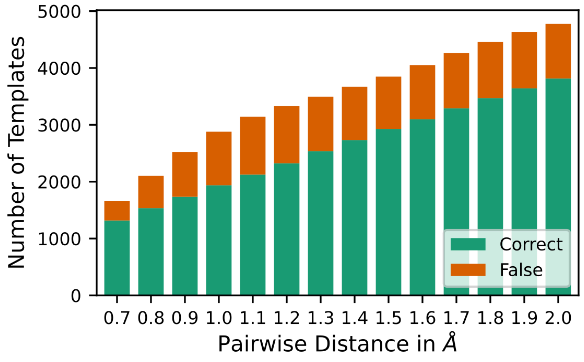


*Figure 4. This bar plot illustrates the total number of templates for which matches were found split by whether at least one match was correct. With increasing pairwise distance, matches to an increasing number of templates were identified. Around 80% of the templates for which any matches were found, produced at least one correct match. However only for about 70% of the templates in our library, were any matches identified in our pdb-validation set.*

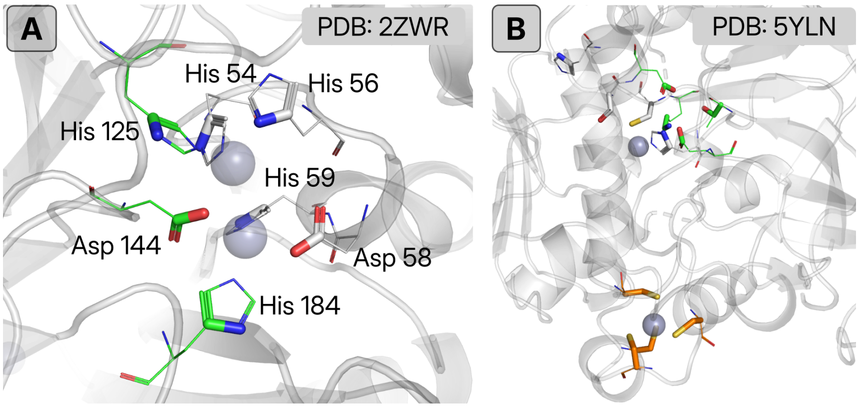


*Figure 5. (A) Annotations for the active site of the bacterial dinuclear zinc beta lactamase (PDB: 2ZWR(2)) could be expanded. The residues His54, His56, Asp58 and His184 are annotated as catalytic (grey residues), while His125, Asp144 and His184 were not (green lines). (B) The bacterial alcohol dehydrogenase 2 (PDB: 5YLN(3)) features two zinc binding sites (grey spheres). One is partially coordinated and located in the active site near both annotated and unannotated catalytic residues (grey and green residues respectively). The other is fully coordinated by four cysteine residues (orange) and has purely structural function. Matched residues predicted as correct are shown as sticks.*

#### Applications of Template Matching

We highlight some examples which illustrate both applications of Enzyme Motif Miner and shortcomings of literature based catalytic annotations such as this. One such application is the annotation of active sites in previously un- or incompletely characterised proteins. Even within our well annotated validation dataset, we observed matches which included unannotated residues in the vicinity of the annotated active site which quite probably contribute to catalysis. By adding such residues to existing annotations of enzymes with matching active sites, Enzyme Motif Miner is suited to expanding annotations for active site residues. Such examples are shown in Fig. S5. Here, the original catalytic annotations for the bacterial dimetallic zinc beta-lactamase B1 (M-CSA entry 15, PDB: 2ZWR*(2)*) His54, His56, Asp58 and His59 could be expanded to include His125, Asp144 and His184 from templates to the Hydroxyacylglutathione hydrolase entry in the M-CSA (entry 157) which also features two coordinated zinc ions in its mechanism.

Templates only account for a tiny fraction of the entire structure which contributes to the function of a protein. In fact, a feature of our template library is the inclusion of templates representing partial active sites which by themselves may only facilitate a proton transfer or metal ion coordination but are insufficient for catalysis. Therefore, matched residues, despite closely resembling an active site, may not be catalytic for a variety of reasons. Since templates do not check for free space or pocket accessibility, matched active sites may be occluded or dysfunctionally altered by additional residues due to charge or steric constraints. In the case of metal binding sites we commonly find matches to residues coordinating structural metal ions. This is illustrated in panel B of Fig. S5 by the bacterial alcohol dehydrogenase 2 (PDB: 5YLN*(3)*). The enzyme has two distinct zinc binding sites. One is catalytically active while the second $Zn^{2+}$ is fully coordinated by four cysteines and only serves a structural purpose. Nonetheless, all coordinating residues for the second ion were matched by our templates. Since the geometry of residues around metal ions is restricted by their possible coordination, such sites are strictly conserved and may have evolved multiple times convergently in different protein families. Without accounting for metal type and full coordination geometry, structural and catalytic metal binding sites are difficult to distinguish. However, while not strictly serving a catalytic purpose, detecting such sites can nonetheless provide contextual insights into protein structure, considering most predicted structures currently lack cofactors.

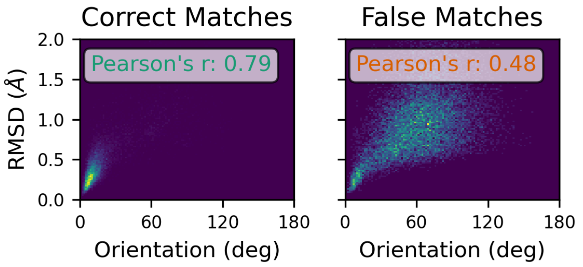


*Figure 6. For correct matches, we found a strong correlation between RMSD and residue orientation. False matches correlated significantly less well. Data is shown for 3-residue matches at 0.8A and 4-residue matches at 1.7A since these datasets are approximately evenly balanced. Pearson's r correlation coefficients are indicated.*

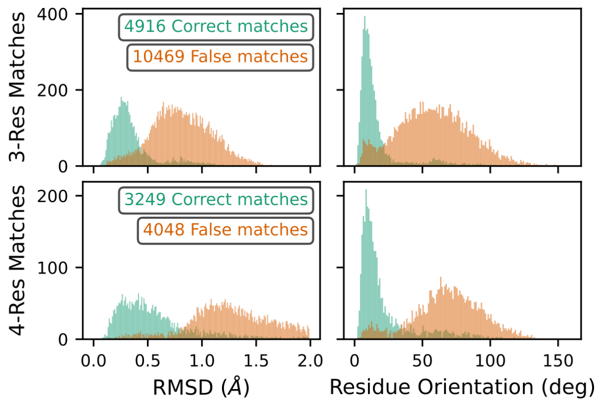


*Figure 7. This histogram illustrates the different distributions of RMSD and residue orientation for 3-residue and 4-residue matches for correct and false matches separately. Data is shown for 3-residue matches at 0.8A and 4-residue matches at 1.7A since these datasets are approximately evenly balanced.*

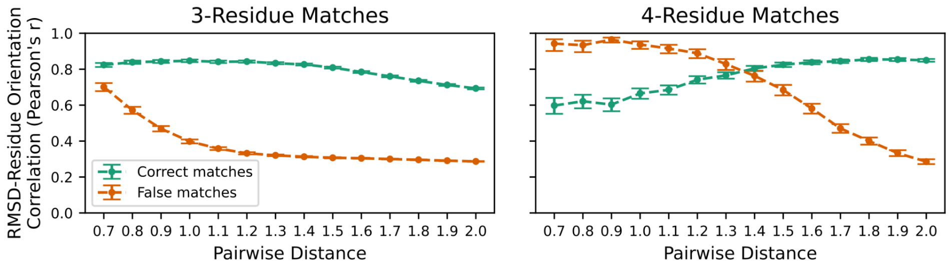


*Figure 8. Pearson's r correlation coefficient with 95% confidence intervals between RMSD and residue orientation for shown separately for correct and false matches with 3- and 4-residues at all pairwise distances.*

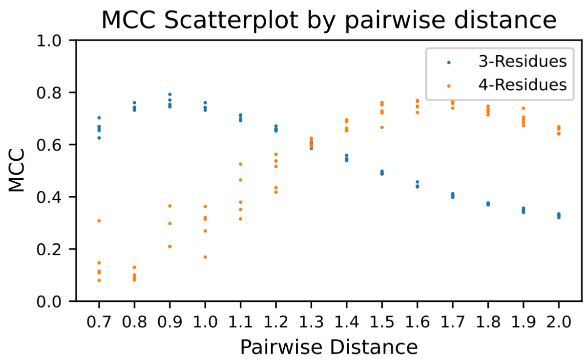


*Figure 9. Matthews Correlation Coefficients for a 5-fold cross-validation of logistic regression models trained to distinguish correct from false matches for matches with 3- or 4-residues at all pairwise distances.*

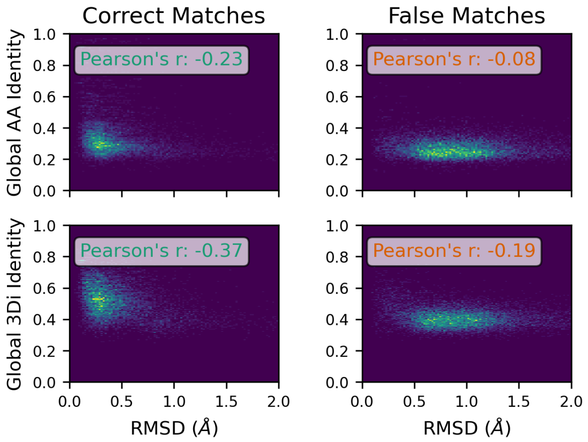


*Figure 10. Only very weak correlation between RMSD and global amino-acid (AA) and foldseek's 3Di identity were found for correct matches. False matches showed no such correlation. Data is shown for 3-residue matches at 0.8A and 4-residue matches at 1.7A since these datasets are approximately evenly balanced. Pearson's r correlation coefficients are indicated. Pearson's r correlation coefficients are indicated.*

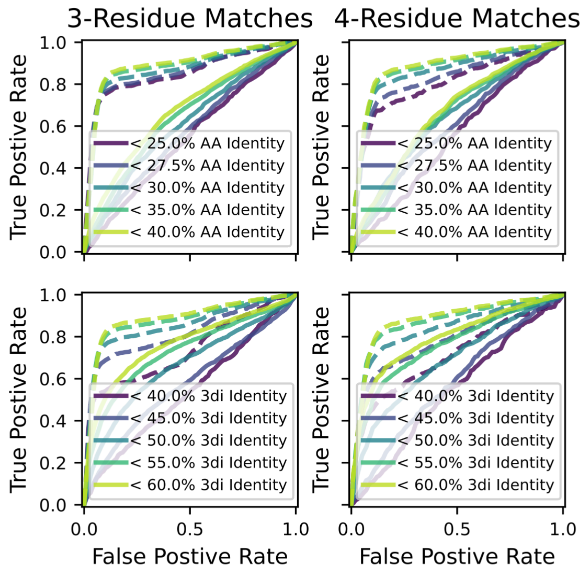


*Figure 11. ROC curves for models trained on RMSD and residue orientation (dashed lines) are compared to models trained on either amino-acid or 3Di sequence identity between template and*

*target (solid lines) to predict correct matches. The relationship between identity and correct matches was further explored by removing match data above a certain maximum sequence identity indicated in the figure legends. Each line represents the mean of a 5-fold cross-validation. Train and test sets were randomly shuffled but stratified to contain similar proportions of correct and false matches. Data is shown for 3-residue matches at 0.8A and 4-residue matches at 1.7A since these datasets are approximately evenly balanced between correct and false matches.*

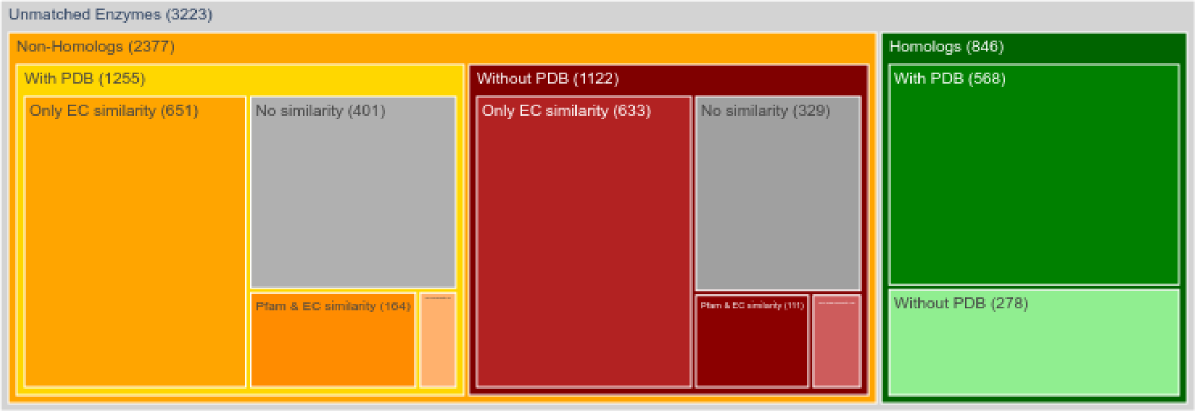


*Figure 12. A tree map illustrating for which unmatched enzymes matches could have been expected based on homology to the M-CSA (green) versus enzymes with no homology (orange and red). Subgroups are colored by resolution in the PDB and further subdivided by EC or Pfam similarity.*

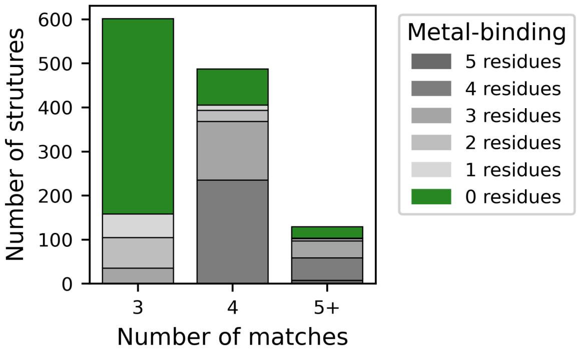


*Figure 13. Matches to Alphafold2 predicted structures of non-enzyme proteins from the SwissProt reviewed human proteome. Of the total 14.875 non-enzyme proteins, 1354 matches were identified to 936 of them. Most of these matches are attributable to metal binding residues. The number of metal binding residues in each match is indicated by color.*

To further analyse matches to Alphafold2 predicted structures of human proteins annotated as non-enzymes by SwissProt (Fig. S13) we manually investigated if any of the 35 5+ residue matches with no metal binding residues might be reasonable. Among others we find a receptor-type tyrosine-protein phosphatase-like N (UniProt: Q16849) for which experiments on the mouse ortholog indicate an aspartic acid outside the catalytic site inhibiting function. Mutating this aspartic acid back to alanine restores catalytic activity*(4)*. A second example is the spliceosome-associated protein CWC27 homolog (UniProt: Q6UX04) for which we find a complete active site but previous studies have deemed it as probable inactive*(5)*.

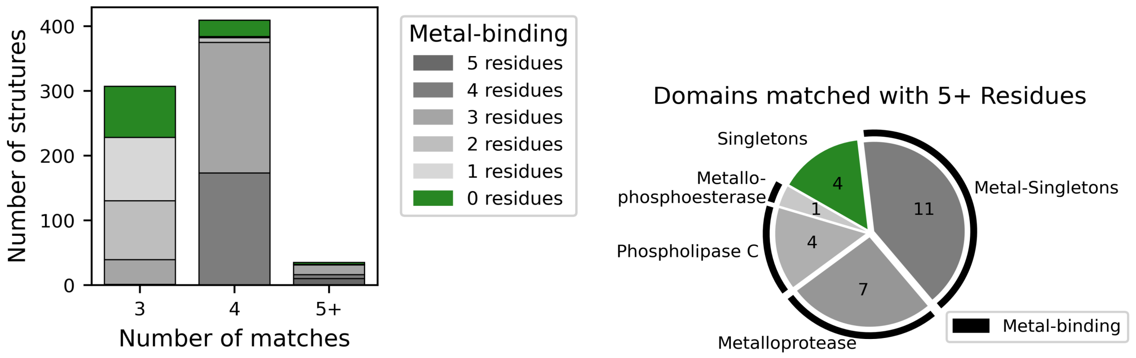


*Figure 14. On 7427 novel folds identified in the encyclopaedia of domains*(6)*, 766 matches to 442 domains were identified. Most of these are attributable to metal binding which is illustrated by the stacked bar plot on the left. 27 domains with 5+ residue matches were more closely analysed. A breakdown is supplied on the right. Only 4 of these domains were matched on non-metal binding sites.*

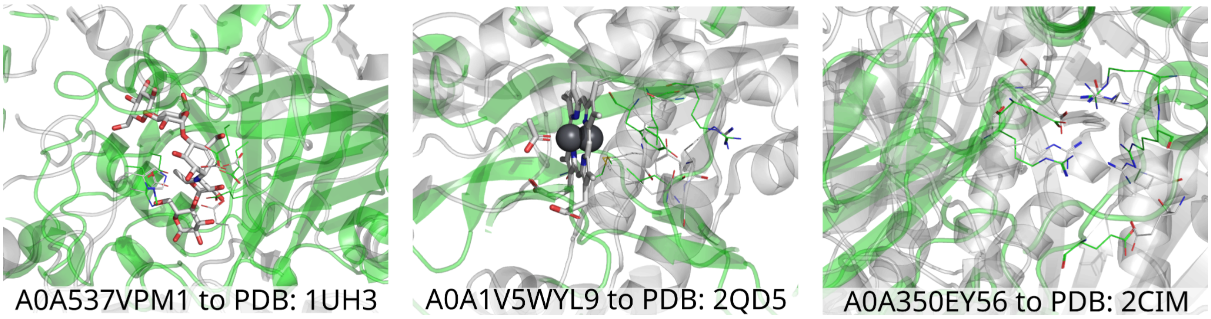


*Figure 15. Novel folds matched by 5+ residue templates at sites which seem at least plausibly catalytic. PDB structures (grey)  including ligands from which the temple was derived are superposed with the novel folds (green) from the encyclopaedia of domains.*

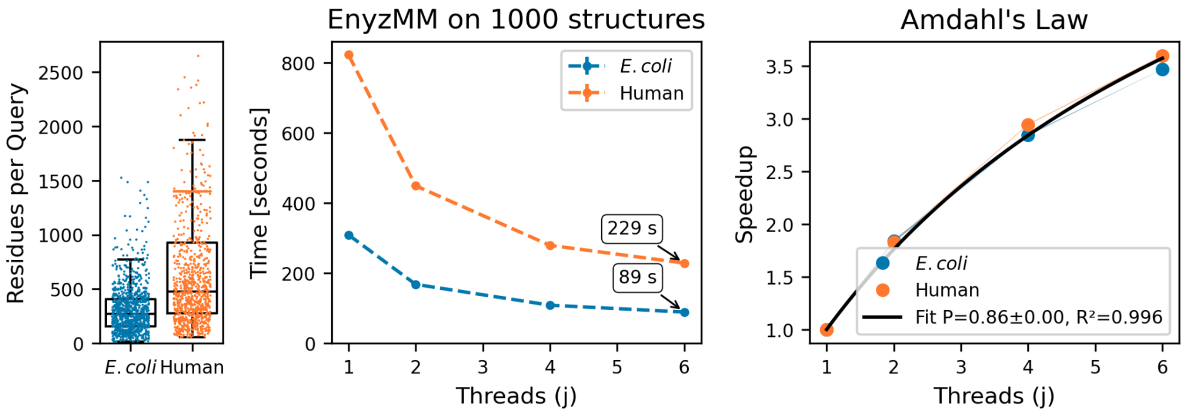


*Figure 16. Performance evaluation of Enzyme Motif Miner on datasets of 1000 E. coli and 1000 human predicted protein structures from the AlphafoldDB. Tests were run on a low-end consumer laptop with an AMD Ryzen 4500U 6-core/6-logical thread CPU on x86_64bit architecture, 8GB RAM and an Ubuntu LTS 24.04 operating system. Tests were run on a warm SSD with default matching parameters while skipping smaller hits. The average E. coli structure has ~250 residues, while the average human structure has ~500 residues. The wall clock time it takes to process the structures is shown by the number of threads used. Fitting the speedup by the number of threads used to Amdahl's law suggests a parallelizable fraction of 86%*(7)*.*
